# Nonlinear super-resolution signal processing allows intracellular tracking of calcium dynamics

**DOI:** 10.1101/2022.01.29.478290

**Authors:** Niccolò Calcini, Angelica da Silva Lantyer, Fleur Zeldenrust, Tansu Celikel

**Affiliations:** Department of Neurophysiology, Donders Institute for Brain, Cognition and Behaviour, Radboud University, Heyedaalseweg 135, Nijmegen, 6525 HJ, The Netherlands; School of Psychology, Georgia Institute of Technology, 654 Cherry Street, Atlanta, GA, 30332, USA

**Keywords:** Calcium imaging, Subcellular localization, Spatiotemporal tracking, Image Series Analysis, Autoregression

## Abstract

Fluorescence imaging of calcium dynamics has revolutionized cellular biology, and especially neuroscience as it allows the study of neural activity across spatially well defined populations. Quantification of fluorescence signals is commonly performed using ratiometric measures. These measures, such as ΔF/F, are easy to implement, but they depend on the definition of a baseline, which is often not trivial outside of the measurement of evoked responses, and needs to be defined for every cell (or other cellular compartment) separately. Here, we present a new quantitative measure of fluorescence by taking advantage of the time dimension of the signal. The new method, which we named ARES (Autoregressive RESiduals), is based on the quantification of residuals after linear autoregression and does not require arbitrary baseline assignment. We detail the basic characteristics of the ARES signal, compare it against ΔF/F, and quantify the improvement in spatial and temporal resolution of recorded calcium data. We further exemplify its utility in the study of intracellular calcium dynamics by describing the propagation of a calcium wave inside a dendrite. As a robust and accurate method for quantifying fluorescence signals, ARES is particularly well-suited for the study of spontaneous as well as evoked calcium dynamics, subcellular localization of calcium signals, and spatiotemporal tracking of calcium dynamics.

**Highlights:**

- A baseline-free quantitative measure of fluorescence is introduced
- The method, called ARES, is based on autoregression
- ARES improves the spatiotemporal resolution of calcium imaging recordings
- Its utility for the localization and spatiotemporal tracking of calcium is shown

## 1. Introduction

Neuronal activity is traditionally studied using either electrophysiological methods or fluorescence, most typically, calcium imaging. Electrical recording techniques provide the best temporal resolution with sampling rates of *>* 20 kHz. However, they rely on inference methods (e.g. spike sorting) to determine cell identities. [1, 2]. Calcium imaging, on the other hand, follows the slower time course of calcium dynamics, at the point of having to rely on an independent set of inference methods to attempt to reconstruct the underlying spiking activity of neurons [3]. However, thanks to the direct photonic observation of networks, calcium imaging enables spatial reconstruction of functional networks [2, 4], including in subcellular resolution [5].

Notwithstanding its importance, calcium imaging has several limitations. First, the nonlinear and slow dynamics of cellular calcium concentrations and indicators hamper the efforts to reconstruct the spiking activity of neurons from calcium traces [3, 6, 7]. Second, chromophores undergo “photobleaching”, where the structure of the molecules themselves is modified during the fluorescence process, making them permanently lose their light emission properties [8]. Photobleaching is particularly prominent during prolonged recording sessions, which is common in functional studies and live recordings of neurons [9]. Light scattering also contributes to the decrease of the spatial signal-to-noise ratio and is often difficult to control [2].

The current standard analysis methods for calcium imaging recordings are ratiometric measures, methods that are based on computing the difference between the signal of interest and a baseline [10], or between two different wavelengths as applied during Fluorescence Resonance Energy Transfer (FRET) imaging [11, 12]. ΔF/F, which is the difference in fluorescence between any frame and a reference (ΔF) in a region of interest (ROI), normalized by the reference value (/F), is by far the most common analysis method used for functional calcium imaging. The simplicity and speed of implementation, and the ease of interpretation of this ratiometric method ensured its widespread implementation. However, ΔF/F has several limitations. A ratiometric measure of this kind has the most efficient application when the baseline remains largely static, otherwise the determination of a correct baseline is non-trivial, requiring for instance kernel density estimates [13], and might greatly affect the outcome of a study. ΔF/F also fails to correct for photobleaching effects in case of long time intervals between the current signal and the baseline value (i.e. prolonged recordings), or unless the baseline is continuously estimated, which is often sub-optimal. Furthermore, ΔF/F does not take advantage of the time dimension present in many applications of the calcium imagine technique which limits its ability to reconstruct the non-linear calcium signals. In an attempt to overcome some of these shortcomings, and to take advantage of the time dimension present in prolonged recordings, we developed ARES (Autoregression RESiduals; https://github.com/DepartmentofNeurophysiology/ARES). ARES is based on the analysis of a sliding time window of fixed length, to which we apply a pixel-wise linear autoregression (predicting a pixel value on the basis of its past values). The time series of the residuals obtained by the comparison between the predicted signal and the measured signal is averaged over a time window to reduce the noise and isolate the transient (novel) events in the signal. Application of ARES to the analysis of calcium dynamics during single-cell stimulation showed that ARES improves the spatial and temporal resolution in functional signals with respect to the raw signal and standard ΔF/F methods. Because ARES provides subcellular information about calcium dynamics, it opens new avenues in high-resolution imaging using calcium indicators.

## 2. Materials and Methods

### 2.1. Experimental methods

All animal procedures were approved by the Radboud University Animal Experiment Committee. Cultures of dissociated cortical neurons were prepared from postnatal day 0 mice on C57BL/6J background. The mother was sacrificed by cervical dislocation, pups by decapitation before their brains were rapidly removed.

#### 2.1.1. Preparation of neocortical dissociated cultures

Brains were put on ice-cold Hank’s balanced salts solution (HBBS; pH 7.3) comprising 10% HBSS without magnesium or calcium (Gibco Life Technologies; Catalog number (Cat.nr.): 14175129), 100 U/ml penicillin-streptomycin (Thermo Fisher Scientific; Cat.nr. 15140122) and 500 *μ*M GlutaMAX™ (Thermo Fisher Scentific; Cat.nr. 35050061). Neocortex was removed bilaterally and incubated in HBSS with 5% trypsin solution (Gibco Life Technologies; Cat.nr. 15090046). The solution was removed and the cells were dissociated using a fire-polished Pasteur’s pipette. Dissociated neurons were plated onto 25 mm coverslips, coated with 1% (weight/volume) poly-L-lysine (Sigma-Aldrich; Cat.nr. 25988-63-0), with a density of 1.6 x 10^5^/coverslip. Cultures were maintained at 37°C in culture medium comprising neurobasal medium (Thermo Fisher Scientific; Catalog number: 21103049) and oxygenated (95%O2/ 5% CO2), supplemented with 2% B27 Supplement (Thermo Fisher Scientific; Cat.nr. 17504044) and 500 *μ*M GlutaMAX™ (Thermo Fisher Scientific; Cat.nr. 35050061). On day 5, the culture medium was replaced with a medium without glutamate. Half the medium was changed twice per week.

#### 2.1.2. Genetically encoded calcium indicator and viral vector production

For visualization of the intracellular calcium dynamics, we virally expressed GCaMP6s [12] under CaMK2 promoter. Viral packaging was adapted from [14, 15, 16]. In short, a suspension of 37.5 *μ*g pRVl (AAV2) (6.25 *μ*g/plate), 37.5 *μ*g pH21 (AAV1) (6.25 *μ*g/plate), 125*μ*g pFΔ6 (Ad Helper) (25 *μ*g/plate), 62.5 *μ*g GCamp6 (12.5 *μ*g/plate), 2 ml CaCl_2_ (2.5 M; Merck; Catalog Cat.nr. 1023820500) and 12 ml of ddH_2_O were slowly added (5 ml/15 cm plate) to HEK293 cells (density: 800.000 cells/plate) as plates were gently swirled. Viral transfection was confirmed 48 hours later with the Eclipse TS100 light microscope (Nikon) coupled with the epi-fluorescence illuminator Nikon Intensilight C-HGFIE and viral particles were purified with heparin (GE Healthcare; Cat.nr. 170406013). The concentration of the virus was checked with qPCR (fwd ITR primer, 5’-GGAACCCCTAGTGATGGAGTT-3’, and rev ITR primer, 5’-CGGCCTCAGTGAGCGA-3’), as in 14. 2 *μ*l of the virus was transferred to dissociated neuronal cell cultures (plate diameter: 25 mm) on day 2 of the culture (DIV2). For these experiments, two batches of the virus with titers of 4.92 x 10^12^ (batch 6) and 1.86 x 10^12^ (batch 7) were used. Transfected neurons were stored in the cell incubator at 37°C until the fluorescence was visible.

#### 2.1.3. Calcium Imaging

Neurons were visualized using LED illumination (Coolled, pE-100) and an exi blue fluorescence microscopy camera (Q-imaging, Model number: EXI-BLU-R-F-M-14-C, CA) coupled to a light microscope (Nikon, FN1) placed on an active vibration isolation table (Table Stable; TS-150). The data was acquired at full field at 10 fps in MicroManager (https://micro-manager.org/) at 14-bit with a readout time of 30 MHz. Data acquisition was triggered using a custom Arduino interface, coupling the whole-cell recording (see below) software Patchmaster (HEKA) with MicroManager, the program controlling the camera and the excitation light source. Electrical recordings, electrical stimulation and calcium imaging were time aligned with clock signals generated in the Patchmaster.

#### 2.1.4. Whole-cell intracellular recording and stimulation

Cell cultures were used 9-15 days after plating, i.e. 7-13 days after viral transfection. Intracellular whole-cell current-clamp recordings were performed as described before [17, 18, 19]. In short, pyramidal neurons were visually selected according to their somatic shape and dendritic morphology. For the duration of the experiments, the culture plate was perfused with Ringer’s solutions (in mM): 10 HEPES (Sigma-Aldrich, Cat.nr. 7364459); 150.1 NaCl; 5 KCl; 1.5 CaCl_2_.2H_2_O; 1 MgCl_2_.6H_2_O; 10 Glucose.H_2_0; pH was adjusted to 7.4 with NaOH (all last chemicals are from Merck, Catalog numbers, respectively are: 7647145, 7447407, 100350408, 1058330250, 14431437). Patch pipette electrodes with a resistance of 5-9 MOhm were pulled from borosilicate glass (Multi Channel Systems; Cat.nr. 300034) using a horizontal puller (Sutter instrument CO. Model P-2000). The intra-pipette solution included (in mM): 5 KCl (Merck, Cat.nr. 7447407); 130 K-Gluconate; 1.5 MgCl_2_.6H_2_O; 0.4 Na3GTP; 4 Na2ATP ; 10 HEPES; 10 Na-phosphocreatine; 0.6 EGTA (Sigma-Aldrich, Catalog numbers, respectively are: 299274, 1058330250, G877, A26209, 7364459, P7936, 67425). pH was set to 7.22 using KOH. (1 M; Merck; Cat.nr. 5033). A chlorided silver wire was used to create electrical continuity between the intra-pipette solution and head-stage, connected to EPC 9 amplifier (HEKA). Current-clamp recordings were performed using step-and-hold pulses, 500 ms in duration. 10 steps of 40 pA current were delivered in every train, and each train was repeated three times. The sweep duration was set to 7 sec. Resting membrane potential was clamped at -70 mV. Evoked calcium dynamics were visualized as described above after binning (x2) with an exposure duration of 100 ms.

### 2.2. Method Description

The main algorithm uses a sliding time window over the frames composing the film in single-pixel resolution. In the following, the time series of a single pixel will be denoted with *P*, and the value of *P* at time (frame) *t* will be denoted with *P*_*t*_. The total number of frames (and the total duration of the time series) is *T* . We compute a linear autoregression of order *k* on the single pixel time series *P* . Calling the *k* parameters of the linear model *U*_*i*_, and the residuals *ϵ*, we have *T* equations and *k* parameters (one equation with *k* parameters for each time point):

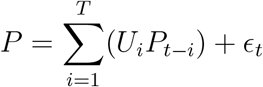

For *k < T*, there are more equations than parameters: this is an over-determined system and it can not be solved exactly. To find the set of parameters *U* = *U*_1_, …, *U*_*k*_ that minimizes the residuals, we use a simple linear least-squares method [20]. Letting *P* be a (*T* × *k*) matrix, having the *i*-lagged time series *P*_*t−i*_ as the *i*-th column, *U* the vector of the *k* model coefficients (*k* × 1) and *E* the vector of the residuals (*T* × 1), we can rewrite the system in matrix multiplication form: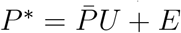. To find the set *Ū* that satisfies

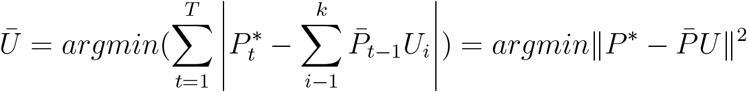

with the assumption that the columns of 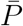 are linearly independent, *Ū* can be computed as:

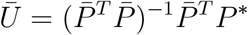

which is solvable by QR decomposition. Having found the coefficients of the linear regression *Ū*, we construct the time series of the residuals *E*(*T*×1) by comparing the measured pixel time series with the ones predicted by the model. Finally, we compute the average over time of the residuals time series *E*, which constitutes the value of a single pixel in an ARES frame:

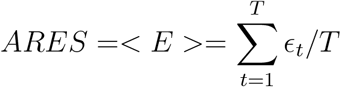

The residuals can be interpreted as noise or unpredictable (novel) events. Thanks to averaging, the noise tends to vanish, especially with longer time windows, while any transient increase or decrease in the signal is still detected. Another way to interpret the average residual time series is to consider that we are using a certain time window (of size *T*) of the signal to predict its future: the less the future time points can be predicted, the more the residuals will differ from zero.

## 3. Results

### 3.1. Performance of ARES relative to ΔF/F

We introduce ARES, a new method for the quantitative analysis of functional calcium imaging data; it does not require any arbitrary assignment of a baseline and deals with the non-linearity in calcium signals. The method is based on a linear prediction of the signal by using its past (autoregression): the prediction error indicating the presence of significant variations, and therefore of an informative signal. This is especially true for calcium signals, as most biological diffusion phenomena, are essentially exponential increases, followed by exponential decays. To avoid considering the error in predicting noise as part of the resulting signal, the prediction errors are averaged over a short temporal window, which contributes to the ‘lag’ between ARES and the raw signal or ΔF/F. This lag decreases with increasing the intensity of the calcium response of a cell (Fig 1 A, C). By using the past of a pixel’s time series to predict its future, we take advantage of the time dimension of the data, including information on the known properties of calcium dynamics (exponential rise and decay), resulting in a better signal to noise ratio (SNR), an improved granularity, and an increased sharpness (amplitude/duration of the signal) with respect to the standard ratiometric normalization ΔF/F methods (Fig 1B, C).

**Figure 1.**
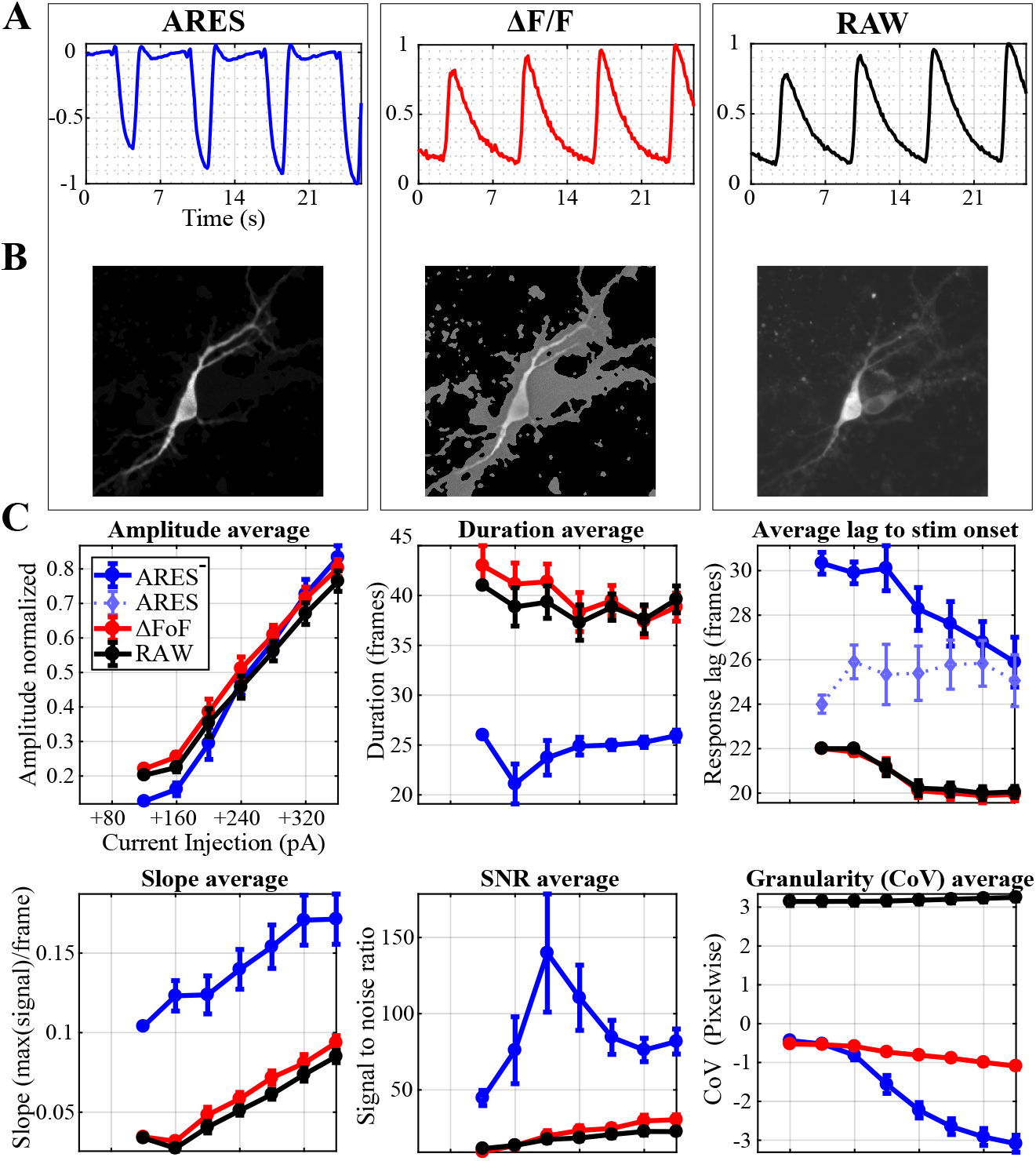
ARES signal and comparison with ΔF/F and raw signals. (A) A current pulse (duration: 500ms; intensity: increasing with increments of +40pA) was injected every 7 seconds. The traces displayed are the average projection on the time axis of each calcium response over the whole region of interest (ROI). Each trace is normalized to the global max in the time series for direct comparisons. (B) Average projection of the absolute value of each signal over the whole time lapse in the ROI. (C) Comparison between ARES, ΔF/F and the raw signal. From top-left to bottom right: 1) the lag between a detectable response in the signal and the stimulus onset (1 frame = 0.1 seconds). 2) The amplitude of each trace, normalize to the maximum signal reached for comparison. 3) The duration of the response to the stimuli provided. 4) The rising slope of the signals. 5) The signal to noise ratio (noise computed by taking a sample of 100 pixels outside the cell soma and dendrites). 6) The coefficient of variation (CoV, measure of granularity of the signal, more negative is a more granular signal). Data shown are the averages and standard errors over 6 cells recordings. Each cell value is, in turn, the average over 3 repetitions of the same stimulation protocol.

### 3.2. Effects of process order and time window

ARES has two main hyper-parameters that need to be determined before the analysis: the order of the autoregressive process used in the autoregression (process order), and the length of the time window on which the analysis is performed (see Method description for details). The first (process order) influences the sensitivity of ARES to shorter or faster dynamics: a smaller process order corresponds to an increased sensibility to shorter transients (Fig 2). The latter, that is the number of consecutive frames taken into consideration in each autoregression (size of the time window), has a similar effect as the process order (Fig 3). Varying the window size has a linear effect on the signal: shortening the time window results in sharper signal and improved SNR, but choosing time windows that are too short (*<*10 frames recorded at 10 Hz) does not allow capturing the entire duration of the calcium transient, hence ultimately reducing the SNR. The parameters can be easily adjusted with the open-source toolbox provided herein (https://github.com/DepartmentofNeurophysiology/ARES).

**Figure 2.**
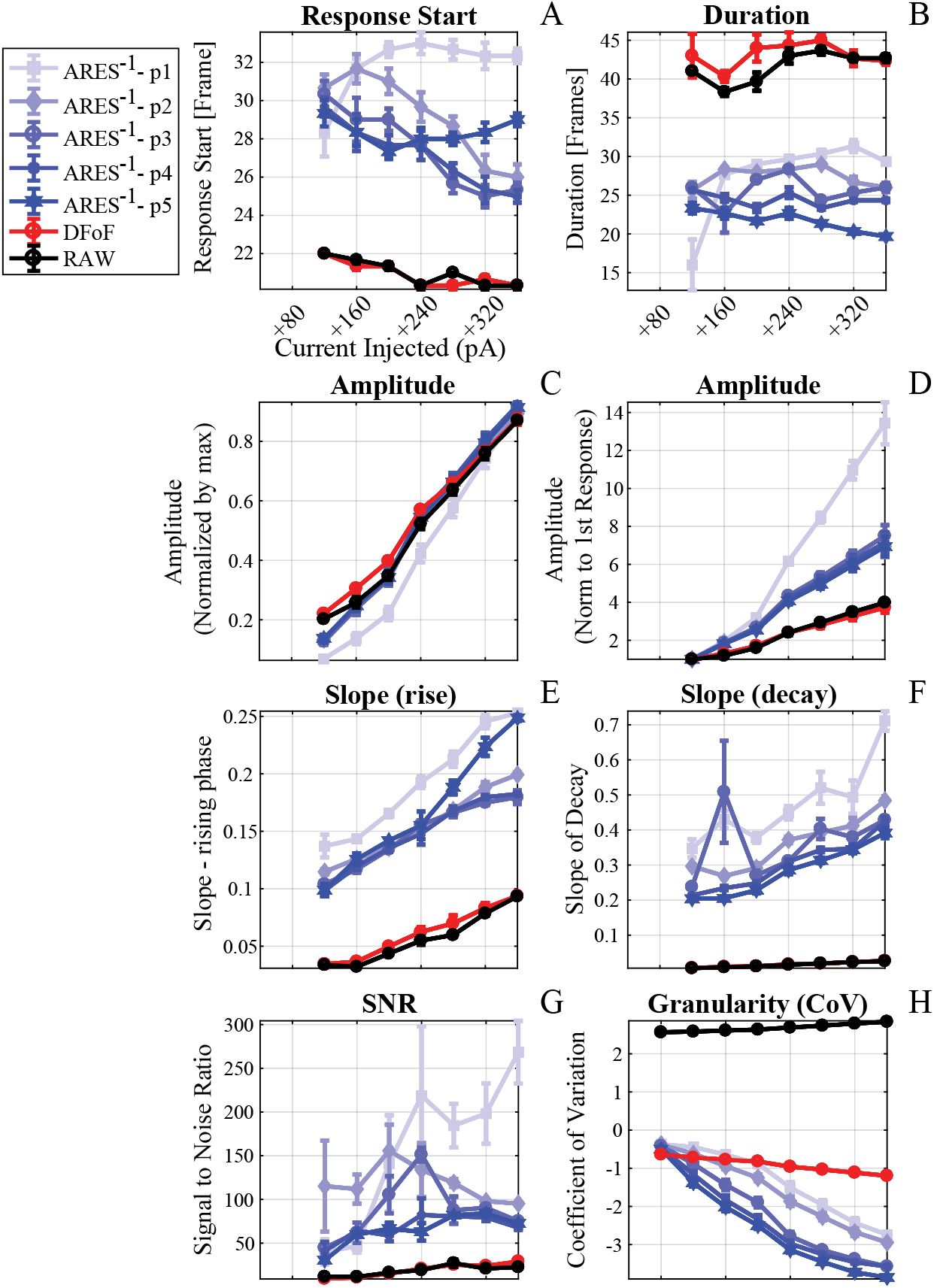
Effects of process order (*p*). The order of the autoregressive process used to predict the signal is a fundamental parameter for the ARES computation. Here we show how its values influence notable characteristics of the output, depending on the stimulation (and therefore response) level, and compare it with the raw signal and ΔF/F. In general, with the exception of the lower possible process order of 1, the results are quite stable and comparable. Most notably, the amplitude of the response (C) from ARES and ΔF/F are the same across all stimulus levels. (A) The number of frames between the stimulus and a detectable response in the signal. (B) The duration in frames of the calcium response. (C) The amplitude of the signal, normalized to the absolute maximum calcium level. (D) The amplitude of the signal normalized to the very first detectable response of the stimuli series. (E) The slope of the rising phase of the signal. (F) The slope of the decay phase. (G) The signal to noise ratio (SNR). (H) The granularity measure of the image. The process order we used by default in other figures was 3.

**Figure 3.**
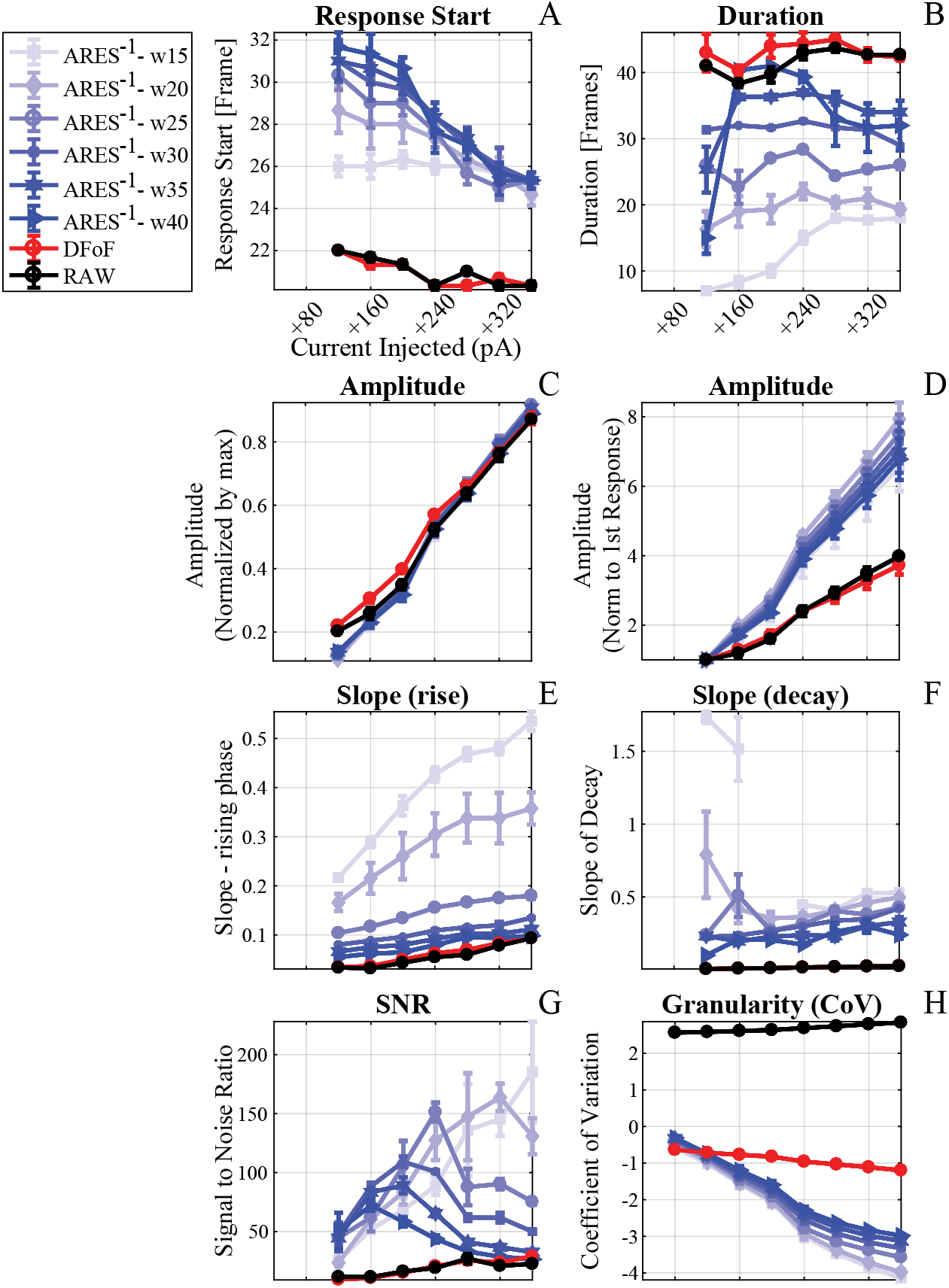
Effects of window size (*w*). The length of the time window considered for the signal prediction is a fundamental parameter for the ARES computation. Here we showed how its length influence notable characteristics of the output, depending on the stimulation level. In general, the effect of varying the window size is mostly linear, with the exception of the amplitude of the response (C) which remains stable across all stimulus levels. (A) The number of frames between the stimulus and a detectable response in the signal. (B) The duration in frames of the calcium response. (C) The amplitude of the signal, normalized to the absolute maximum calcium level. (D) The amplitude of the signal normalized to the very first detectable response of the stimuli series. (E) The slope of the rising phase of the signal. (F) The slope of the decay phase. (G) The signal to noise ratio (SNR). (H) The granularity measure of the image. The window length we used in our other examples by default in our figures was 25 frames.

### 3.3. Tracking wavefronts

Spatial projections of ARES (see Fig 1B) suggest that ARES also provides high-resolution spatial information about the source of the signal. Therefore we tested the algorithm for tracking calcium wavefronts. In-vitro bright field calcium imaging data were collected in primary neuron cultures as described (see Materials and Methods).

ARES presents an increased SNR and at the same time a signal that maintains the same amplitude of the calcium signal and the ΔF/F response, but without a very long tail decay (Fig 1). This allows us to observe and measure the intracellular calcium influx in the form of a propagating wave coming from a dendritic spine (Fig 4 and Supplementary Video 1). Unlike ARES, ΔF/F is not able to resolve how the propagation of the wavefront along the dendrite.

**Figure 4.**
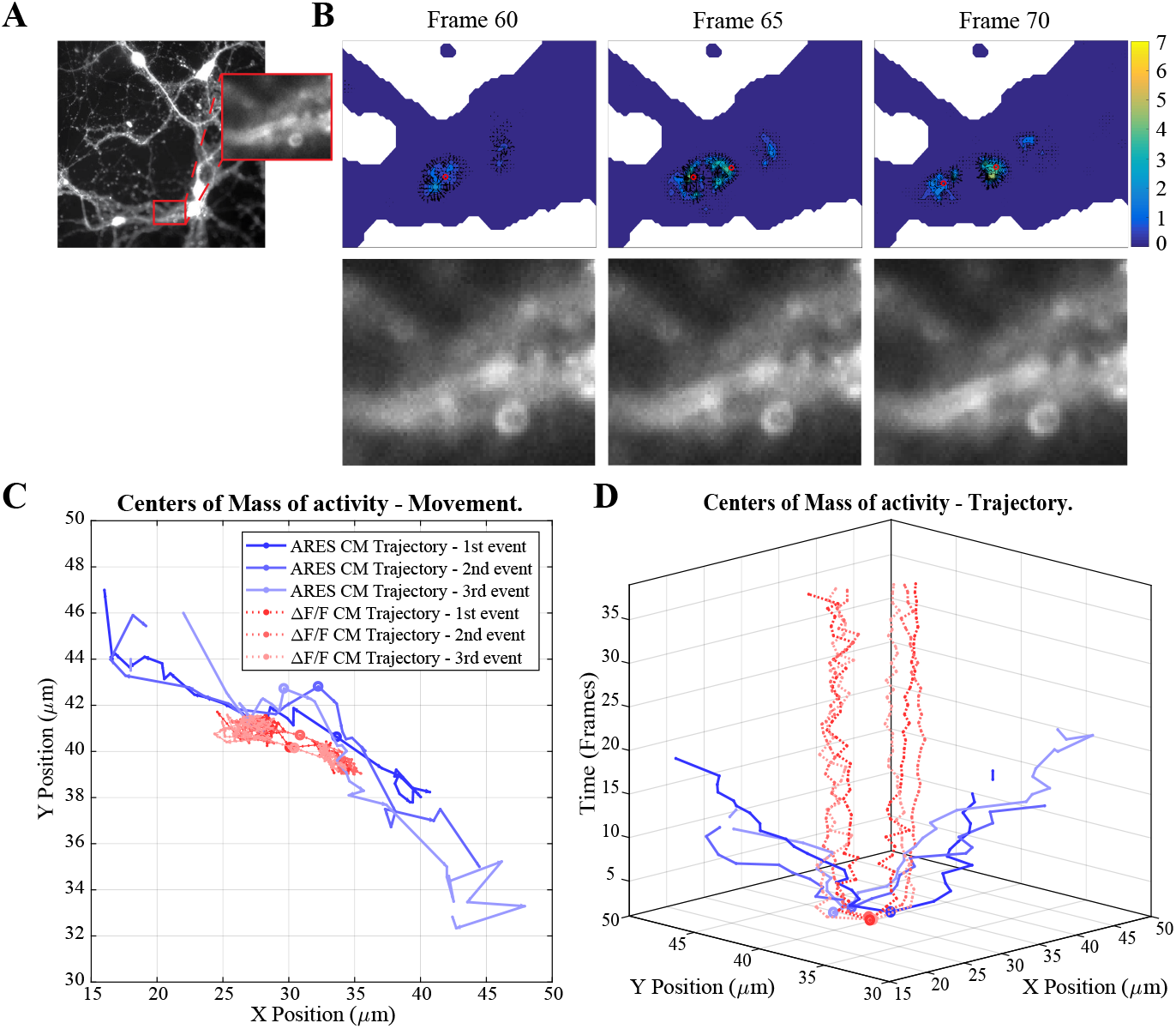
Spatial analysis of subcellular calcium dynamics with ARES. (A) The ROI in a dendritic spine. (B) The propagation of a calcium wave in consecutive frames (see Supplemental Film 2). Top row: the color code represents ARES values. The red circles indicate the position of the center of mass of the activity (the wavefront). Bottom row: corresponding images analyzed with ΔF/F. (C) The 2-D movement tracking of the wave front, tracked as the position of their center of mass, and (D) its trajectory together with time axis. Different shading of the same color represent different events recorded from the same spine. In comparison to ARES, ΔF/F (red) is unable to pick up any dynamic of the wave propagation.

## 4. Discussion

Ratiometric methods such as ΔF/F are intrinsically limited by the necessity of a comparison with a single baseline value. We herein introduced a method, called ARES, that extrapolates the temporal dynamics of the signal in a pixel-wise fashion, and uses this knowledge to detect transient events. We showed that the method improves the temporal and spatial resolution of calcium imaging signals, while maintaining the same linearity properties as ΔF/F (Fig 1C). As an example application, we showed that ARES allows for the study of intracellular calcium dynamics, in the form of wave propagation from spines into dendrites.

The ARES signal is characterized by small positive increments in the signal, representing sudden rises in the calcium signal, and by larger negative troughs, representing the beginning of the signal decay. Once the decay in calcium concentration becomes linearly predictable, ARES takes a value of zero as there is no longer any difference between the predicted and actual signal trace. Linear regressions are not able to correctly capture the dynamics of calcium, which are defined by exponential rises or decays, but as the linear prediction fails whenever there is a novel calcium event, the non-linear dynamics are captured in the residuals. In considering the residuals themselves, instead of the regression coefficients, anything that is not fitted by the linear regression (including the calcium transients) is now represented in the processed signal.

The improved signal resolution and transient detection of ARES, together with its baseline independence and time locality, make the method ideal for the detection of small and non-stimulus locked events, and for studying their dynamic properties. Furthermore, the toolbox accompanying the method provides an intuitive user interface for the use with in-vitro recordings, and an explicit example of the application of the method. The super-resolution imaging of signal transients might offer other interesting advantages such as the differentiation between intra- and extra-cellular calcium sources, and a better reconstruction of the spiking activity, which remains a major challenge in the field.[6, 7].

## Supplementary Material

Supplemental Video 1. **ARES allows the tracking of a dendritic calcium wave front** The video shows imaging of a dendrite taken from primary neuronal cultures (see Material and Methods) at 10 Hz. Four videos are presented side by side for comparison: the raw signal, a relative heatmap, and the heatmaps of ΔF/F and the positive part of the ARES analyzed signal, with tracking of the wavefront displayed as a small red dot.

## Acknowledgments

We thank Dr. Eric Jansen for help with the cell culture preparation, and the members of the Department of Neurophysiology for stimulating discussions. This work was supported by grants from the European Commission (Horizon2020, nr. 660328), European Regional Development Fund (MIND, nr. 122035) and the Netherlands Organisation for Scientific Research (NWO-ALW Open Competition, nr. 824.14.022) to TC, the Netherlands Organisation for Scientific Research (NWO-Veni, nr. 863.150.25) to FZ, and a doctoral fellowship by the National Council for Scientific and Technological Development of Brazil (CNPQ) to ASL.

